# Arginine methylation coordinates with phosphorylation to regulate NADP^+^ synthesis

**DOI:** 10.1101/2024.10.11.617635

**Authors:** Yue Zhu, Wanjun Zhang, Mengxue Hu, Xiaoke Xing, Yanze Li, Hemeng Zhang, Yuanyuan Zhao, Luojun Chen, Chunmin Zhang, Yali Chen, Weijie Qin, Suxia Han, Huadong Pei, Pingfeng Zhang

## Abstract

NADK is the sole cytosolic enzyme responsible for synthesizing NADP^+^ from NAD^+^ within cells. The homeostasis of NAD^+^/NADP^+^ is controlled by NADK, usually dysregulated in various cancers, yet the precise underlying regulatory mechanisms remain largely unknown. In this study, we discover that PRMT6 methylates NADK at R39, R41, and R45, resulting in the suppression of NADK kinase activity and NADP^+^ synthesis. Mutations of these sites promote cancer cell proliferation and tumor growth. PRMT6-mediated methylation of NADK coordinates with Akt-mediated phosphorylation to regulate NADK, which stimulates its activity to increase NADP^+^ production through relief of an autoinhibitory function inherent to its amino terminus. PRMT6 also inhibits NADK activity in a phosphorylation-independent manner. Furthermore, NADK methylation is upregulated by RB1/E2F pathway in high-fat diet mice. Downregulation of NADK methylation in HCC enhances NADP^+^ production to promote cancer development. Our findings illuminate the molecular regulatory mechanisms governing NAD^+^/NADP^+^ homeostasis, suggesting that the PRMT6-NADK axis emerges as a direct player in high energy state. Furthermore, our research suggests potential therapeutic targets through chemical and genetic interventions.

**Significance:** Previous studies revealed that phosphorylation of NADK activates its activity in response to nutrient or other signaling pathways. Here, we discovered that PRMT6 methylates NADK at residues R39, R41, and R45, leading to the suppression of NADK kinase activity and NADP^+^ synthesis. This PRMT6-mediated methylation works in conjunction with Akt-mediated phosphorylation to regulate NADK. Additionally, we found that NADK methylation is upregulated by the RB1/E2F pathway in mice on a high-fat diet. In hepatocellular carcinoma (HCC), the downregulation of NADK methylation enhances NADP^+^ production, thereby promoting cancer development. Our findings provide a significant breakthrough in understanding the regulation of NADK and its response to environmental factors.

## Introduction

The NAD^+^/reduced NAD^+^ (NADH) and NADP^+^/reduced NADP^+^ (NADPH) redox couples play key roles in maintaining cellular redox homeostasis and modulating numerous biological events, including cell metabolism and growth ^1–3^. Approximately 10% of cellular NAD^+^ is phosphorylated by NAD^+^ kinases into NADP^+4,5^, which can be dephosphorylated to NAD^+^ by NADP^+^ phosphatases ^6–9^. The NAD^+^/NADP^+^ ratio is modulated by many physiological conditions, including nutrient availability, hormone levels, and oxygen levels, among others. However, the underlying molecular mechanisms driving these regulatory processes remain incompletely understood. Deficiency or imbalance of NAD^+^/NADP^+^ has been associated with many pathological disorders, such as aging and cancers ^5,10,11^. NADK is the sole cytosolic enzyme that catalyzes the phosphorylation of NAD^+^ to form NADP^+^, thus precise modulation of NADK is vital for cell survival and growth ^12,13^. A recent study showed that phosphoinositide 3-kinase (PI3K)-Akt pathway directly stimulates NADP^+^ synthesis through Akt-mediated phosphorylation of NADK. Akt phosphorylates NADK on three serine residues (Ser44, Ser46, and Ser48) within an amino-terminal domain. This phosphorylation stimulates NADK activity to increase NADP^+^ production through relief of an autoinhibitory function inherent to its amino terminus^14^.

Arginine methylation is a ubiquitous posttranslational modification (PTM) that regulates critical cellular processes, including signal transduction, DNA replication, and gene transcription, among others^15^. Aberrant methylation of arginine residues is a common PTM increasingly associated with cancer, obesity, and other human diseases^16^. There are three forms of arginine methylation, including monomethylated arginine (MMA), symmetrical dimethylated arginine (sDMA), and asymmetrical demethylated arginine (aDMA). Post-arginine monomethylation (Rme1) serves as an intermediate, and type II protein arginine methyltransferases (PRMTs) generate symmetric arginine dimethylation (sDMA), while type I PRMTs, including PRMT6, catalyze asymmetric arginine dimethylation (aDMA)^17^. An increasing body of literature suggests that PRMT6 may have tumor-suppressive or tumor-promoting roles in different types of cancers^18^. However, the role of PRMT6 in NAD^+^ metabolism and in HCC remains largely unknown.

In this study, we uncovered novel regulatory mechanisms governing NADK in response to nutrient availability in HCC. We found that PRMT6 methylates NADK, the sole enzyme for NADP^+^ production from NAD^+^, and PRMT6-mediated NADK methylation suppresses NADK kinase activity and NADP^+^ production, at least partially through antagonizing Akt-mediated phosphorylation. Moreover, the PRMT6-NADK axis emerges as a direct player in HCC development.

## Results

### The arginine methylation of NADK at the N-terminal region suppresses NADP^+^ production

NAD^+^ kinase plays a key role in regulating NAD^+^/NADP^+^ homeostasis, thereby exerting control over cellular catabolic and anabolic metabolism. Previous research revealed that NADK undergoes phosphorylation at its N-terminal regulatory region to upregulate its activity in response to nutrient stimulation^14,19,20^. As cells adapt to diverse physiological conditions, NADK may employ different mechanisms in response to various stimuli. This prompted us to investigate whether there are additional regulatory pathways or PTMs modulating NADK activity in HCC. Endogenous NADK protein was purified from HEK293T cells using anti-NADK antibody, then subjected to protease digestion and mass-spectrometry analysis. In addition to the previously reported phosphorylation modifications (Fig. S1A, B), NADK was mono-methylated and di-methylated at Arg39, Arg41, and Arg45 (Fig. 1A). Mono-methylation of NADK was detected by anti-mono-methyl arginine antibody, but di-methylation could not be detected by anti-di-Methyl arginine antibody (Fig. S1C). We generated NADK R39K, R41K, R45K, and a R3K mutant (R39K/R41K/R45K) with all three sites mutated to lysine (Fig. S1D). The three individual mutants had reduced mono-methylation, and the mono-methylation of R3K mutant was totally abolished (Fig. S1D). These data indicate that Arg39, Arg41, and Arg45 are the main methylation sites on NADK. The Arg39, Arg41 and Arg45 residues in human NADK are conserved throughout vertebrates (Fig. 1B).

**Figure 1.**
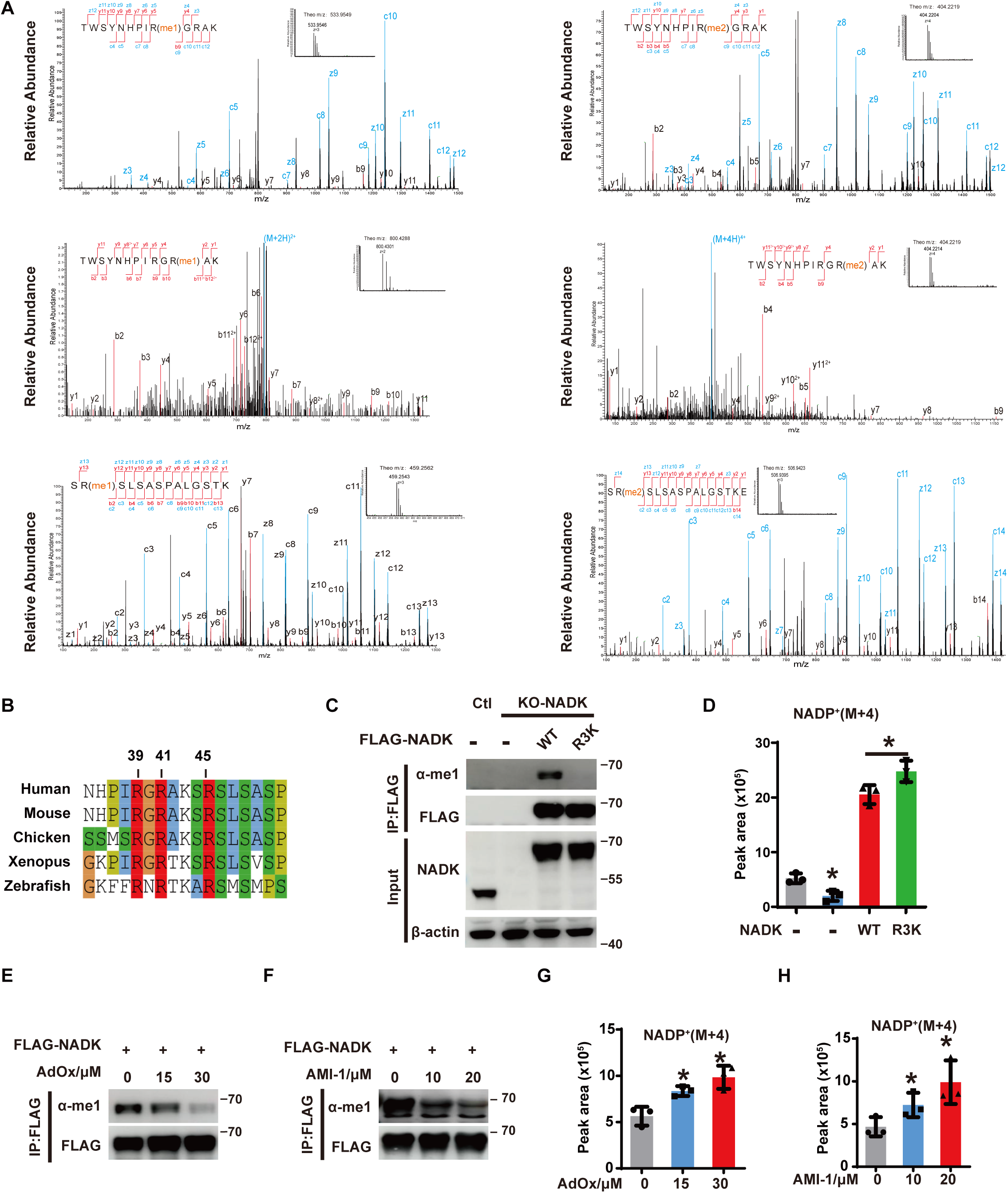
Arginine methylation of NADK at N-terminal region suppresses NADP^+^ production. (A) Representative MS/MS spectrum for peptides containing mono- or di-methylation on residues Arg39, Arg41, Arg45. Endogenous NADK was immunopurified from HEK293T cells and analyzed by LC-MS/MS as described in the methods. (B) Sequence alignment of the N-terminal region that is involved in methylation in NADK. Three arginine residues with mono-/di-methylation are highlighted in red. (C) Parental or NADK-KO HEK293T cells were transfected with the indicated plasmids, and the methylation of NADK WT or R3K mutant was detected by immunoprecipitation and western blotting with an anti-Mono-Methyl Arginine antibody. (D) As in (C), parental or NADK-depleted HEK293T cells were reconstituted with NADK WT or R3K mutant. *De novo* NADP^+^ synthesis were quantified of by LC-MS. (E-F) Global methylation inhibitors AdOx (E) and AMI-1 (F) was applied in a dose gradient to inhibit NADK methylation in HEK293T cells. (G-H) Quantification of *de novo* NADP^+^ synthesis by LC-MS in HEK293T cells when treated with methylation inhibitors AdOx (G) and AMI-1 (H). [(D), (G) and (H)] Data are presented as the mean ± SD of biological triplicates and are representative of at least three independent experiments. *P < 0.05 for pairwise comparisons calculated using a two-tailed Student’s t test.

To determine the functional significance of NADK methylation, endogenous NADK was depleted in HEK293T cells, followed by stable reconstitution with wild-type NADK and R3K mutant. The exogenous proteins were expressed at similar levels but had elevated abundance compared to endogenous NADK (Fig. 1C). The R3K mutant abolished NADK methylation (Fig. 1C). NADP^+^ is synthesized from NAD^+^ phosphorylation by NADK, or re-oxidized from NADPH via several anabolic pathways^21^. To investigate the impact of NADK methylation on NADP^+^ production from NAD^+^ phosphorylation, we employed stable isotope tracing with ^13^C3-^15^N-nicotinamide using a 1-hour pulse label, as previously reported^14^ (Fig. S1E). Inhibition of the NAD^+^ salvage enzyme nicotinamide phosphoribosyltransferase (NAMPT) with the specific inhibitor FK866^22^ decreased the appearance and fractional enrichment of the M+4 derivatives of NAD^+^ and NADP^+^, serving as pathway validation (Fig. S1F). ^13^C3-^15^N-nicotinamide tracing demonstrated that the R3K mutant exhibited an accelerated ability to synthesize NADP^+^ compared to that of the wild-type NADK (Fig. 1D). These data demonstrate that NADK methylation on R39/R41/R45 inhibits NADP^+^ synthesis.

Consistent with the above results, global methylation inhibitors AdOx^23^ and AMI1^24^ also inhibited NADK methylation in a dose-dependent manner and promoted *de novo*-synthesized NADP^+^ production (Fig. 1E-H).

### PRMT6 interacts with and methylates NADK *in vitro* and in cells

Arginine methylation catalyzed by PRMT is a common PTM in mammalian cells, and PRMTs target RG-rich and non-RG-rich motifs with similar frequency to control protein stability and function^25,26^. In mammals, there are 9 protein arginine methyltransferases (PRMT1-9) ^16,27,28^. PRMT1 and PRMT6 were found to contribute to NADK di-methylation, as overexpression of PRMT1 or PRMT6 decreased the mono-methylation of NADK, resulting in more conversion to di-methylation (Fig. 2A). This is consistent with previous literature^29,30^. We also performed an *in vitro* methylation assay using purified NADK and PRMT1/PRMT6 from *E. coli*. Both PRMT1 and PRMT6 were able to methylate NADK *in vitro*, with PRMT6 showing stronger activity (Fig. 2B). Knockdown of PRMT6 with siRNA in cells significantly decreased the mono-methylation of NADK (Fig. S2A).

**Figure 2.**
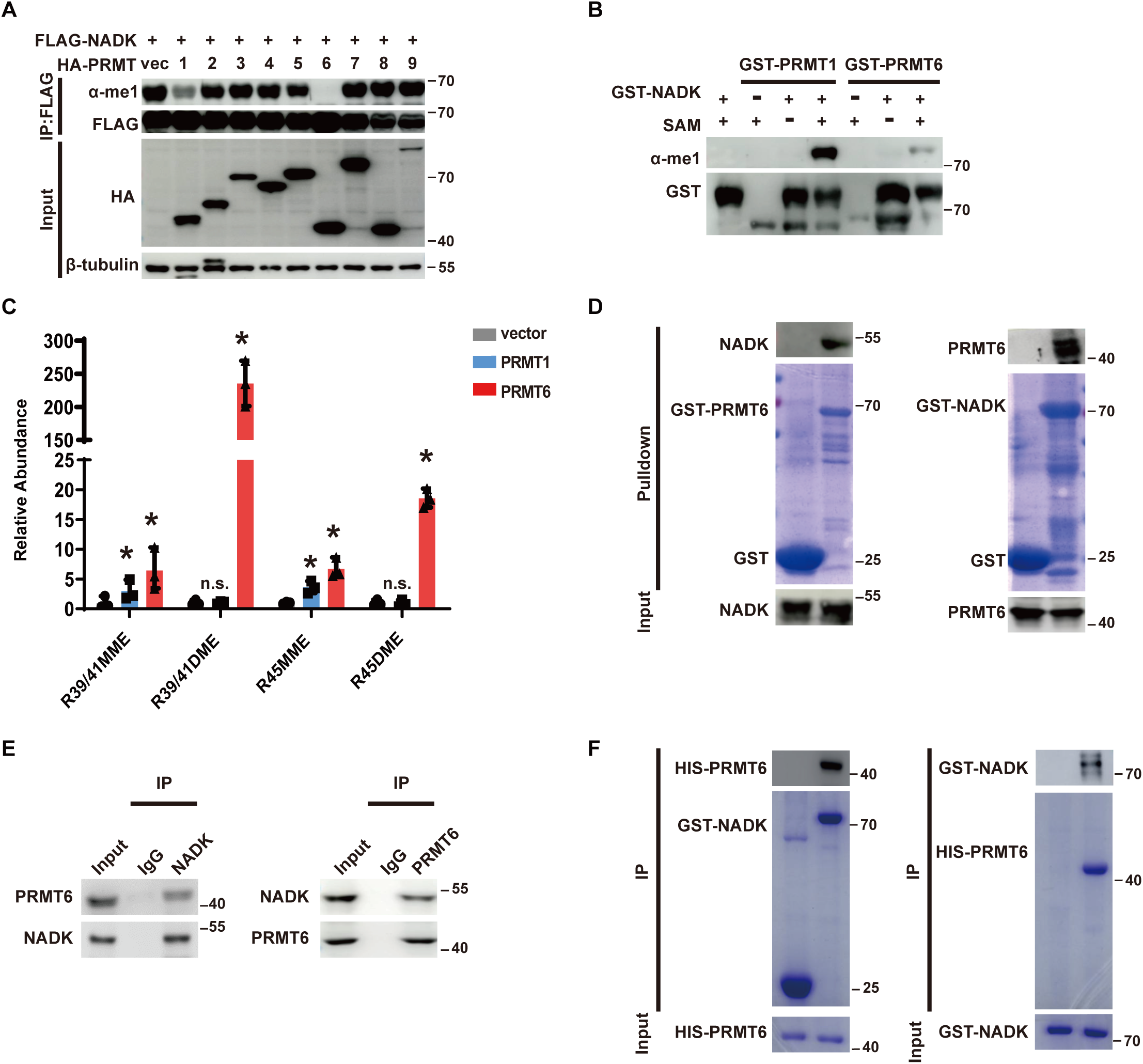
PRMT6 methylates NADK *in vitro* and in cells to inhibit its activity. (A) NADK methylation assay was performed with FLAG-NADK co-transfected with HA-PRMT1-9 into HEK293T cells, respectively. NADK methylation were detected with an anti-Mono-Methyl Arginine antibody. (B) *In vitro* NADK methylation assay was performed with GST-NADK as a substrate. Purified GST-PRMT1 and PRMT6 were used in the assay, and NADK methylation were detected as in (A). (C) *In vivo* NADK methylation quantification was performed with PRMT1 or PRMT6 overexpressed in HEK293T cells. FLAG-NADK was immunopurified first and then subjected to LC-MS/MS analysis. Mono-/di-methylation on NADK R39/41 and R45 were analyzed separately. (D) GST pull-down assays were performed to detect the interactions between NADK and PRMT6 *in vitro*. All GST proteins were produced from *E. coli*. Left: purified GST-PRMT6 pulled down NADK overexpressed in HEK293T cells. Right: purified GST-NADK pulled down PRMT6 overexpressed in HEK293T cells. (E) Reciprocal endogenous immunoprecipitation (IP) was performed to detect interactions between NADK and PRMT6 in HEK293T cells. (F) *In vitro* protein-protein interaction assays were performed to detect the direct interactions between NADK and PRMT6 with purified proteins. All proteins were overexpressed and purified from *E. coli*. Left: purified GST- NADK pulled down HIS-PRMT6. Right: purified HIS-PRMT6 pulled down GST-NADK. [(C)] Data are presented as the mean ± SD of biological triplicates and are representative of at least three independent experiments. *P < 0.05 for pairwise comparisons calculated using a two-tailed Student’s t test.

Moreover, mass spectrometry-based LFQ was applied to evaluate NADK methylation by PRMTs. PRMT1 and PRMT6 were overexpressed in HEK293T cells, then NADK proteins were purified and the methylation levels were analyzed with LC-MS/MS method. R39/41 and R45 di-methylation increased dramatically when PRMT6 was overexpressed. Meanwhile, mono-methylation on R39/41 and R45 also increased a few times (Fig. 2C). However, PRMT1 overexpression only slightly increase mono-methylation, but had no significant effect on NADK di-methylation in cells (Fig. 2C). Consistent with these results, overexpression of wild type PRMT6, but not the enzyme deficient mutant, deceased the mono-methylation of wild type NADK but not the NADK R3K mutant (Fig. S2B, C).

Moreover, GST pull-down and co-IP assays supported the direct interaction between PRMT6 and NADK *in vitro* and in cells (Fig. 2D-F). Furthermore, we mapped the regions of PRMT6 and NADK responsible for this interaction by expressing truncated PRMT6 or NADK mutants in HEK293T cells, followed by a co-IP assay (Fig. S2D, E). The D1 truncation mutant of PRMT6, which contains the N-terminal Rossmann Fold (1-182) and lacks the β barrel domain and dimerization helices, (residues 183–375), abolished the binding with NADK, indicating that the interacting region is located in C-terminal domain. The D2 and D3 constructs, which contain the conserved catalytic domain (86-182) or a variable N-terminal region (1-85) to the C-terminal domain, respectively, both showed similar but reduced interaction with NADK, suggesting that they play a similar helper role in the interaction with NADK (Fig. S2D). Similarly, we generated truncation mutants of NADK (Fig. S2E). The PRMT6-binding region of NADK was mapped to the C-terminal catalytic domain (N2 construct, residues 106–446) (Fig. S2E). Notably, comparison of WT and N2 construct shows that the N-terminal region (residues 1-105), which contains the substrate residues of PRMT6, enhances the interaction between NADK and PRMT6 (Fig. S2E). All these data support that PRMT6 is the major methyltransferase for NADK.

### PRMT6-mediated NADK methylation attenuates its kinase activity

Next, we explored whether and how PRMT6 influences NADK activity. Accurately quantification using synthetic unmodified and methylated standard NADK peptides showed that overexpression of PRMT6 in HEK293T cells increases the NADK methylation 2∼4 times (Fig. 3A, B), and it inhibits *de novo* NADP^+^ synthesis in a PRMT6 dose-dependent manner (Figs. 3C and EV3A). The newly synthesized NADP^+^ partially represents the NADP^+^ synthesis rate. To understand how NADK methylation affects the NAD^+^/NADP^+^ homeostasis in cells, we also measured the total NAD^+^ and NADP^+^ levels in HepG2 cells. Accompanying an increase in NAD^+^ level, the total NADP^+^ level decreased (Fig. S3B). Accordingly, NADPH level also reduced, while GSH level remains unchanged (Fig. S3C, D), indicating NADP^+^ to NADPH is more essential to cell homeostasis. On the other hand, knockdown of PRMT6 with siRNA promotes NADP^+^ production and the total NADP^+^ level (Figs. 3D and EV3E, F). The PRMT6-specific inhibitor EPZ020411^31,32^ had a similar effect (Figs. 3E and EV3G, H). These results indicate that PRMT6 inhibits *de novo* NADP^+^ production, thus has an negative effect on total NADP^+^ level.

**Figure 3.**
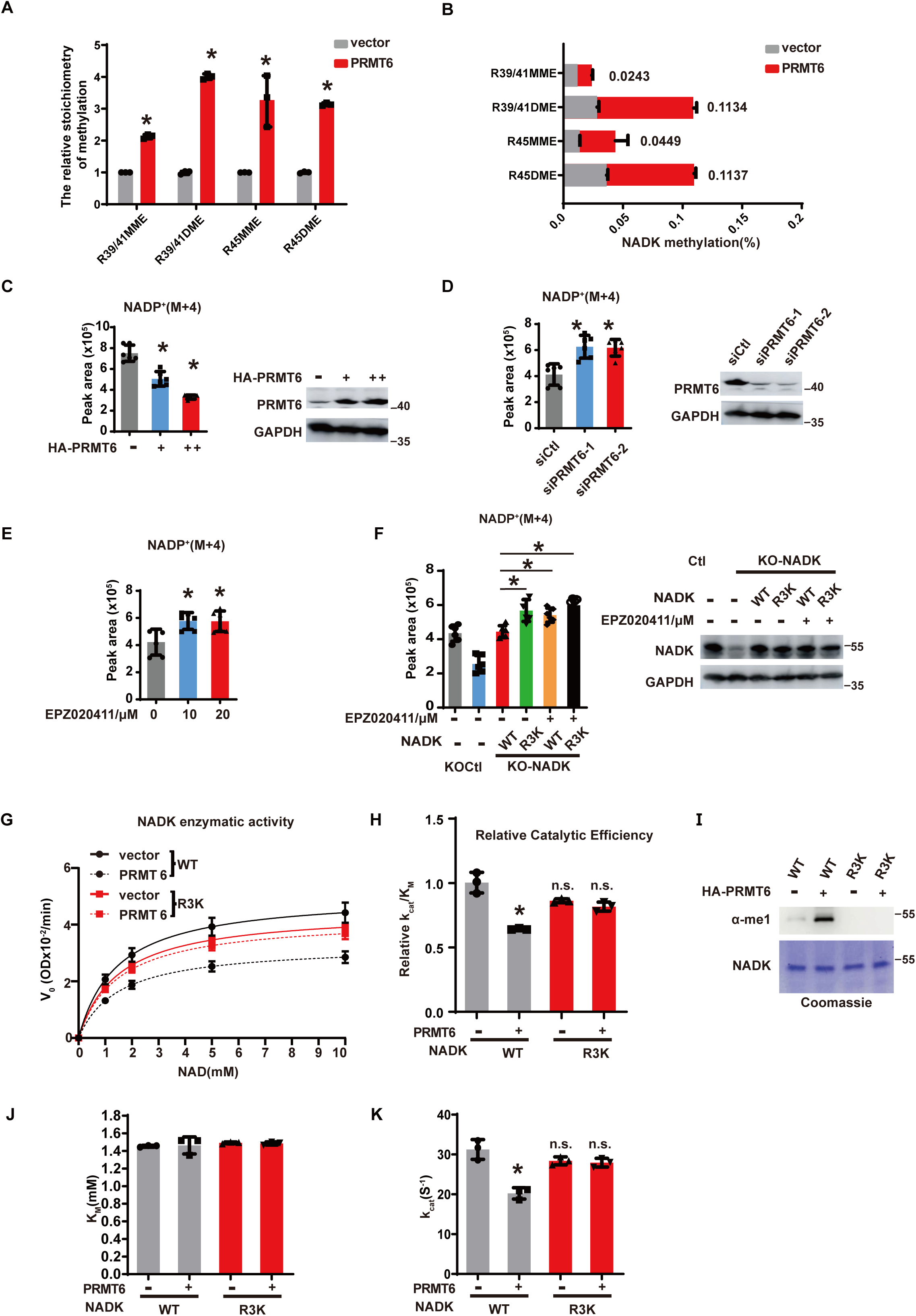
PRMT6-mediated NADK methylation attenuates its kinase activity. (A-B) *In vivo* NADK methylation quantification was performed with or without PRMT6 overexpression in HEK293T cells. NADK was immunopurified first and then subjected to LC-MS/MS analysis. Mono-/di-methylation on NADK R39/41 and R45 were analyzed using synthetic unmodified and methylated NADK standard peptides. (A) represents the quantitative values of NADK methylation when comparing PRMT6 overexpression to non-overexpression, while (B) depicts the proportions of methylated NADK among the total NADK peptides. (C-D) Quantification of *de novo* NADP^+^ synthesis by LC-MS. HepG2 cells were transfected with HA-PRMT6 plasmids (C, dose dependent) or siRNAs (D). PRMT6 protein levels were detected by western blot with indicated antibodies. (E) HepG2 cells were treated with the PRMT6-specific inhibitor EPZ020411 in a dose gradient to inhibit NADK methylation. (F) Quantification of *de novo* NADP^+^ synthesis by LC-MS. NADK-depleted HepG2 cells were reconstituted with NADK WT or R3K mutant and treated with the PRMT6-specific inhibitor EPZ020411. NADK methylation was detected with anti-Mono-Methyl Arginine antibody. (G-I) NADK activity assay for NADK WT or R3K mutant. NADK proteins were immunopurified from HEK293T cells and subjected to *in vitro* methylation assay with purified PRMT6 as described in methods. Reaction velocity (G), relative catalytic efficiency (H), NADK protein and methylation quantification (I), KM (J), Kcat (K) are presented. [(A-F] Data are shown as the mean ± SD (n=6). [(G-K)] Data are presented as the mean ± SD of biological triplicates and are representative of at least three independent experiments. *P < 0.05 for pairwise comparisons calculated using a two-tailed Student’s t test.

We went further to check the PRMT6 inhibitor EPZ020411 on NADP^+^ production. EPZ020411 does not change the NAD^+^ level, however, it significantly increases the newly synthesized NADP^+^ and the total NADP^+^ levels. Moreover, NADK R3K mutant cells had more *de novo* synthesized NADP^+^ compared to that of wild-type cells. While PRMT6 inhibitor increases NADP^+^ production on WT NADK, it has no effect on the R3K mutant (Figs. 3F and EV3I). Overexpression of PRMT6 inhibits NADP^+^ production in wild-type cells but not in NADK R3K mutant cells (Fig. S3J), indicating that PRMT6 inhibits NDAP^+^ synthesis through methylation of NADK on R39/R41/R45. The *in vitro* NADK activity assay revealed that PRMT6-mediated methylation decreased the activity of wild-type NADK, but had no effect on the activity of the R3K mutant of NADK (Fig. 3G, H). Direct *in vitro* methylation of NADK by PRMT6 also decreased its catalytic efficiency (Fig. 3I-K). These data demonstrate that PRMT6 inhibits the synthesis of NADP^+^ due to direct inhibitory effects on NADK enzymatic activity through methylation of R39/R41/R45 within its N-terminal region.

To investigate how NADK methylation affects its activity, we examined the subcellular localization, stability, and tetrameric organization of NADK. PRMT6 does not affect NADK subcellular localization in the cell (Fig. S4A, B), and PRMT6-mediated methylation also does not affect NADK stability (Fig. S4C). Additionally, NADK methylation does not affect its tetrameric organization (unpublished data). To further explore how NADK methylation affects its kinetics, NADK was methylated by PRMT6 *in vitro* and purified for the measurement of NAD^+^ and ATP dissociation constant via Isothermal Titration Calorimetry (ITC) assay. Compared to the non-methylated NADK, methylated NADK has slightly lower affinity to NAD^+^, and the affinity to ATP is significantly impaired (Fig. S4D-G), partially explains its lower activity. Taken together, all these data suggest that N-terminal methylation slightly affects its binding affinity to its substrates, perhaps by enhancing an autoinhibitory function inherent to its amino terminus.

### NADK methylation crosstalks with its phosphorylation

Akt phosphorylates NADK on Ser44, Ser46, and Ser48, relieves the N-terminal autoinhibition and stimulates NADK activity ^14,20^. We found that overexpression of PRMT6 abolished NADK phosphorylation on Ser44, Ser46, and Ser48 (Fig. 4A, B), indicating that PRMT6-mediated NADK methylation inhibits its phosphorylation. On the other hand, Akt-induced phosphorylation of NADK diminished its methylation (Fig. 4C). Thus, methylation and phosphorylation occur in the same regulatory N-terminal region of NADK and antagonize each other.

**Figure 4.**
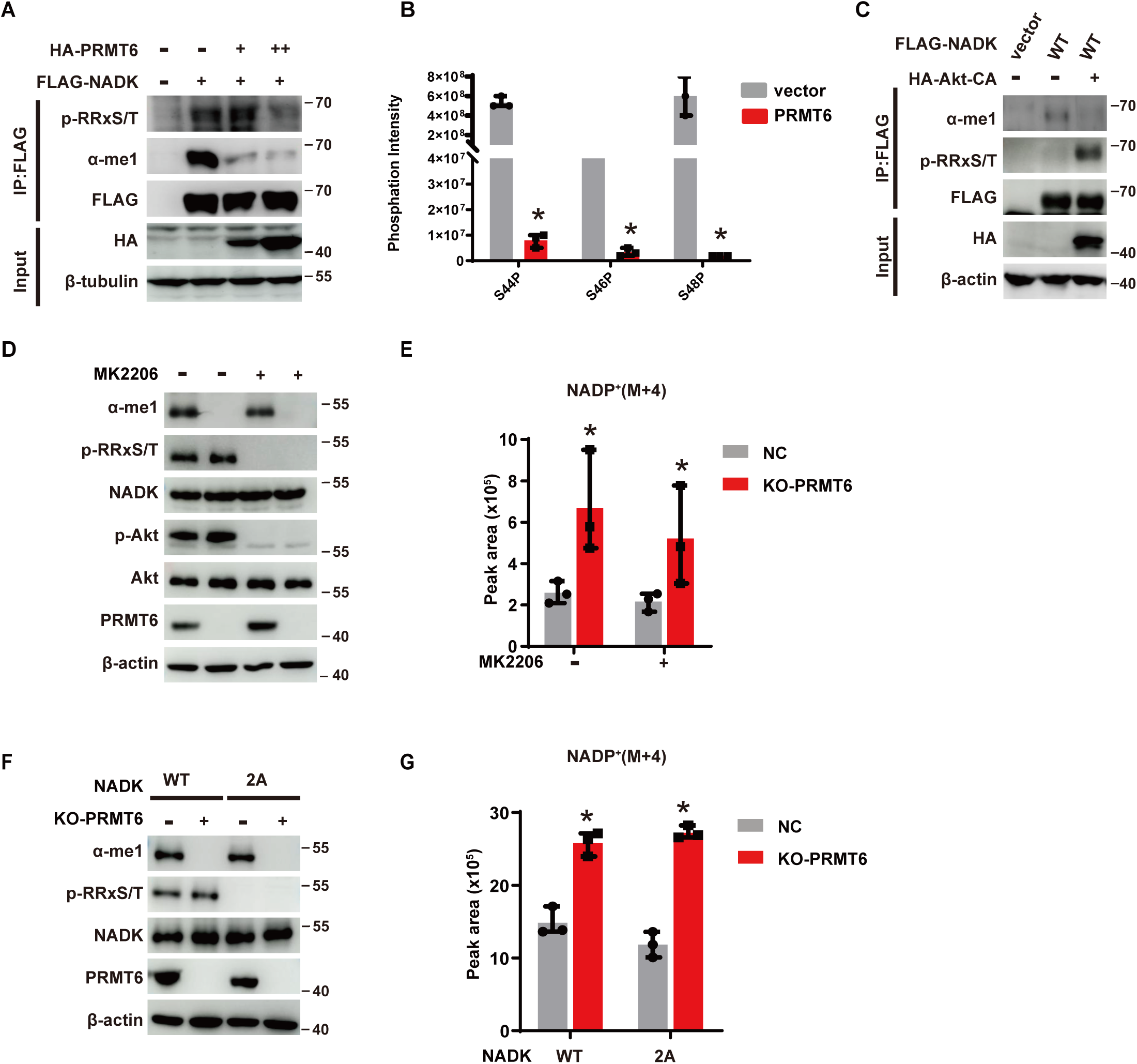
NADK methylation crosstalks with its phosphorylation. (A) HEK293T cells were co-transfected with FLAG-NADK and HA-PRMT6. FLAG immunoprecipitates were immunoblotted with the indicated antibodies to detect NADK methylation and phosphorylation. (B) *In vivo* NADK phosphorylation quantification was performed with or without PRMT6 overexpression in HEK293T cells. NADK was immunopurified first and then subjected to LC-MS/MS analysis. (C) As in (A), but cells were co-transfected with HA-Akt-CA instead of PRMT6. (D-E) PRMT6 was knocked out in HEK293T cells and treated with Akt inhibitor MK2206. The NADK protein phosphorylation levels and its methylation levels were quantified with indicated antibodies (D). Quantification of *de novo* NADP^+^ synthesis by LC-MS were presented in (E). (F-G) NADK WT or 2A mutant were overexpressed in HEK293T cells with or without PRMT6 knockout. The NADK protein phosphorylation level and its methylation level were quantified with indicated antibodies (F). Quantification of *de novo* NADP^+^ synthesis by LC-MS were presented in (G). [(B), (E) and (G)] Data are presented as the mean ± SD of biological triplicates and are representative of at least three independent experiments. *P < 0.05 for pairwise comparisons calculated using a two-tailed Student’s t test.

We further investigated if PRMT6 inhibits NADK activity in a Akt-mediated phosphorylation-dependent manner. Akt inhibitor MK2206 suppresses NADP^+^ production independent of PRMT6 expression (Fig. 4D, E). PRMT6 knockout promotes NADP^+^ production in both wild-type cells and NADK phosphorylation-deficient cells (S44/46A, 2A) to the same extent, and there is no significant difference between these two cell types (Fig. 4F, G). Therefore, we conclude that PRMT6-mediated methylation inhibits NADK activity at least partially through antagonizing Akt-mediated phosphorylation in wild-type cells, but PRMT6 also inhibits NADK activity in Akt-mediated phosphorylation-deficient cells by directly enhancing the autoinhibition conformation. Structural details of this autoinhibitory mechanism needs further study.

### Arginine methylation of NADK suppresses cancer cell proliferation

NADK is a promising target for cancer treatment, as cancer cells often exhibit an elevated NADPH pool to support their rapid proliferation^21,33^. Thus we explored whether NADK methylation has an inhibitory effect on HCC cell proliferation. As shown in Fig. 5A-C, we generated NADK knockout HepG2 cancer cells by using CRISPR/Cas9 and reconstituted these cells with wild-type NADK or a methylation-deficient mutant NADK (R3K). We found that NADK knockout dramatically inhibited cancer cell proliferation and colony formation, and wild-type NADK was able to rescue this effect; interestingly, the NADK methylation-deficient mutant further boosted cancer cell proliferation and colony formation (Fig. 5A-C).

**Figure 5.**
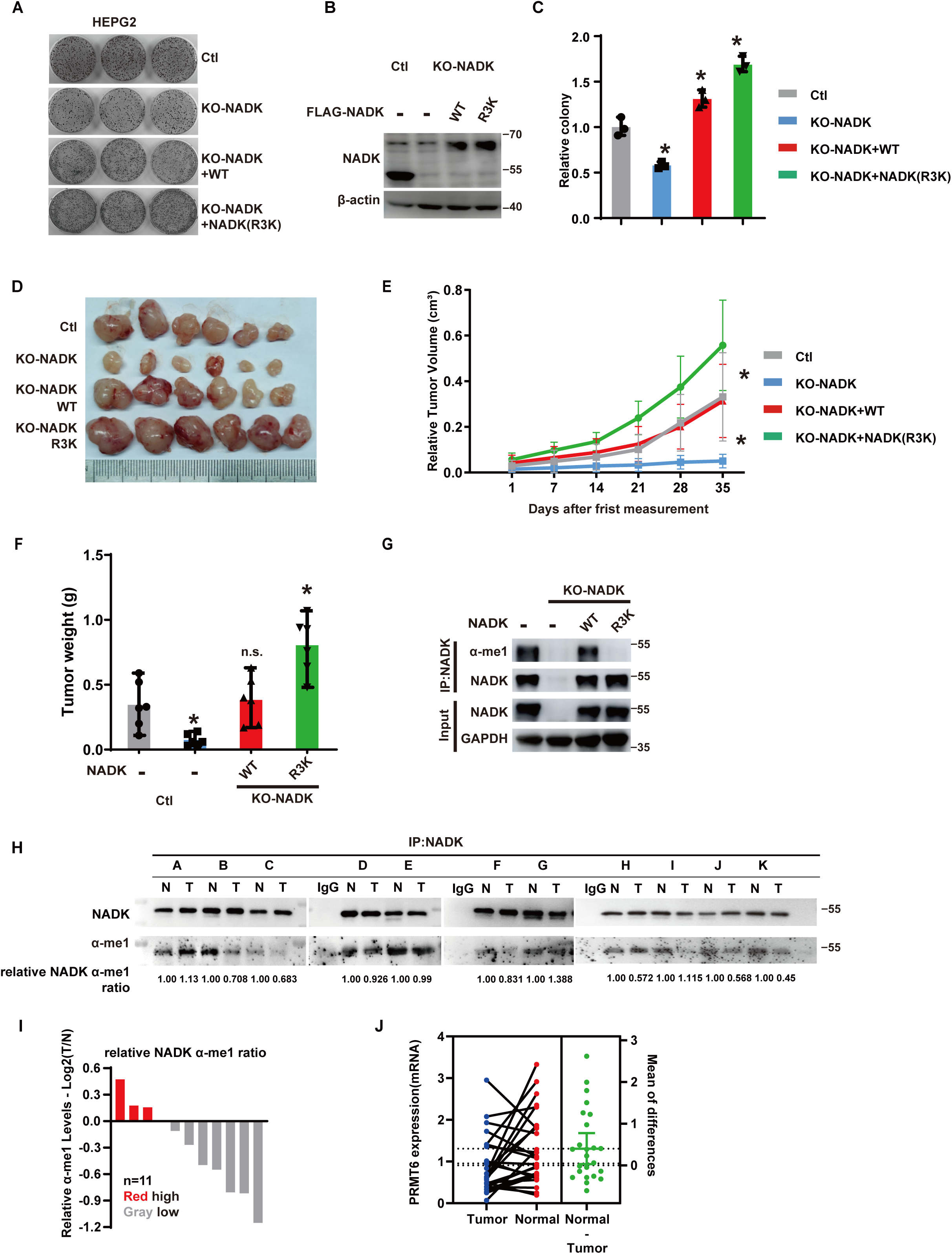
NADK methylation suppresses tumorigenesis. (A-C) NADK-depleted HepG2 cells were transfected with FLAG-NADK WT and R3K mutant plasmids, and the resulting colonies were fixed and stained with crystal violet (A). NADK expression level was detected with anti-NADK antibody (B). Relative colony numbers were shown in (C). (D-G) PRMT6-mediated methylation of NADK suppresses tumor growth in xenografted mice. Tumors were collected after mice were sacrificed at the endpoint. Tumor images (D), tumor volume (E) and tumor weight (F) were presented (n=6). NADK expression and methylation levels in the tumor were detected by western blot with indicated antibodies (G). (H-I) Endogenous NADK proteins were immunoprecipitated from liver cancer tissues and patient-matched normal liver tissues. The NADK methylation were detected via immunoblotting using the indicated antibodies (H). The relative NADK methylation ratio was determined by arbitrarily setting the α-me1/NADK to 1 in the normal tissue, and the relative α-me1/NADK in the tumor was calculated accordingly. The intensity of the bands was analyzed by ImageJ software (I). (J) mRNA levels of PRMT6 expression were assessed in liver cancer tissues and patient-matched normal liver tissues (n=25). [(C)] Data are presented as the mean ± SD of biological triplicates and are representative of at least three independent experiments. [(E) and (F)] Data are shown as the mean ± SD (n=6). *P < 0.05 for pairwise comparisons calculated using a two-tailed Student’s t test.

To test the *in vivo* effect of NADK methylation, HepG2 cells reconstituted with NADK WT and R3K mutant were injected into BALB/c nude mice. While NADK knockout cells significantly inhibit tumor growth, WT NADK rescues the tumor growth, and the R3K mutant further stimulates tumor growth (Fig. 5D-G). Collectively, these data support a model in which NADK methylation inhibits cancer cell proliferation.

### Downregulation of NADK methylation in HCC enhances NADP^+^ production to promote cancer development

PRMT6 plays different roles in various cancers, it has been found to be downregulated and to play suppressive roles in ovarian cancer, breast cancer, prostate cancer, melanoma, and liver cancer^18^. To mimic the PRMT6-mediated NADK methylation deficiency in liver cancer, the NADK depleted liver cancer cell line HepG2 was reconstituted with the R3K NADK mutant and then injected into obese BALB/c nude mice. The tumors were harvested to measure the NADK methylation, NADP^+^ production, and metabolic phonotype. Indeed, while several serum metabolites show obese phenotype (Fig. S5A-E), the R3K mutant diminishes the NADK methylation (Fig. S5F), and significantly promotes tumor growth (Fig. S5G-I) via enhancing NADP^+^ production (Fig. S5J). In the meanwhile, most metabolites in the tumor, including nucleotide metabolites, glucose metabolites, and amino acids metabolites (Fig. S5K-M) remained unchanged, or showed little to no steady-state changes, highlighting the suppressing role of PRMT6-NADK-NADP^+^ pathway in the tumor development.

We further investigated NADK methylation in liver cancer patients. PRMT6 is frequently downregulated in tumor compared to the patient-matched normal tissues (mRNA levels), and correspondingly, NADK methylation levels were decreased (Fig. 5H-J), indicating reduced PRMT6-mediated NADK methylation in liver tumors. This finding is consistent to previous reports on the downregulation of PRMT6 in HCC^34,35^. Given the tumor diversity and heterogeneity, discriminating the PRMT6-NADK pathway in liver cancer patients may help improve cancer treatment.

### Upregulation of the PRMT6-NADK axis inhibits NADP^+^ synthesis in high-fat diet mice

Previous literature has shown altered NAD^+^ metabolism in metabolic disorders. So we further created a high-fat diet (HFD) induced obese mice model to study how high-energy physiological conditions affect NAD^+^ metabolism. The body weight of high-fat diet induced obese mice was significantly increased compared to the mice fed with a standard chow diet (Fig. S6A, B). We also monitored the metabolic changes after HFD intervention, the glucose, total cholesterol (TCHO), triglycerol (TG), high density lipoprotein (HDL), and low density lipoprotein (LDL) all increased in HFD mice, showing an obese phenotype (Fig. S6C-G). As most NAD^+^ is biosynthesized in the liver^34^, we checked NAD^+^/NADP^+^ metabolism and the NADK activity in the liver. The liver volume of HFD obese mice was significantly increased (Fig. S6H), and the H&E staining of liver sections showed significant lipid droplets (Fig. S6I). NAD^+^/NADH levels didn’t change too much, but NADP^+^/NADPH was significantly reduced in HFD obese mice (Fig. 6A, B), which is also consistent with a recent study ^35^. While NADK protein level was unchanged in the liver of HFD obese mice (Fig. 6C), PRMT6 mRNA level and protein level increased significantly (Fig. 6C, D). Mass spectrometry and western blot results showed the Arg45 di-methylation also increased (Fig. 6C, E), which suggests the NADK activity is inhibited by elevated methylation. These results characterize a new negative regulatory pathway for NADK via PRMT6-mediated NADK methylation, induced by high-fat diet or in obesity.

**Figure 6.**
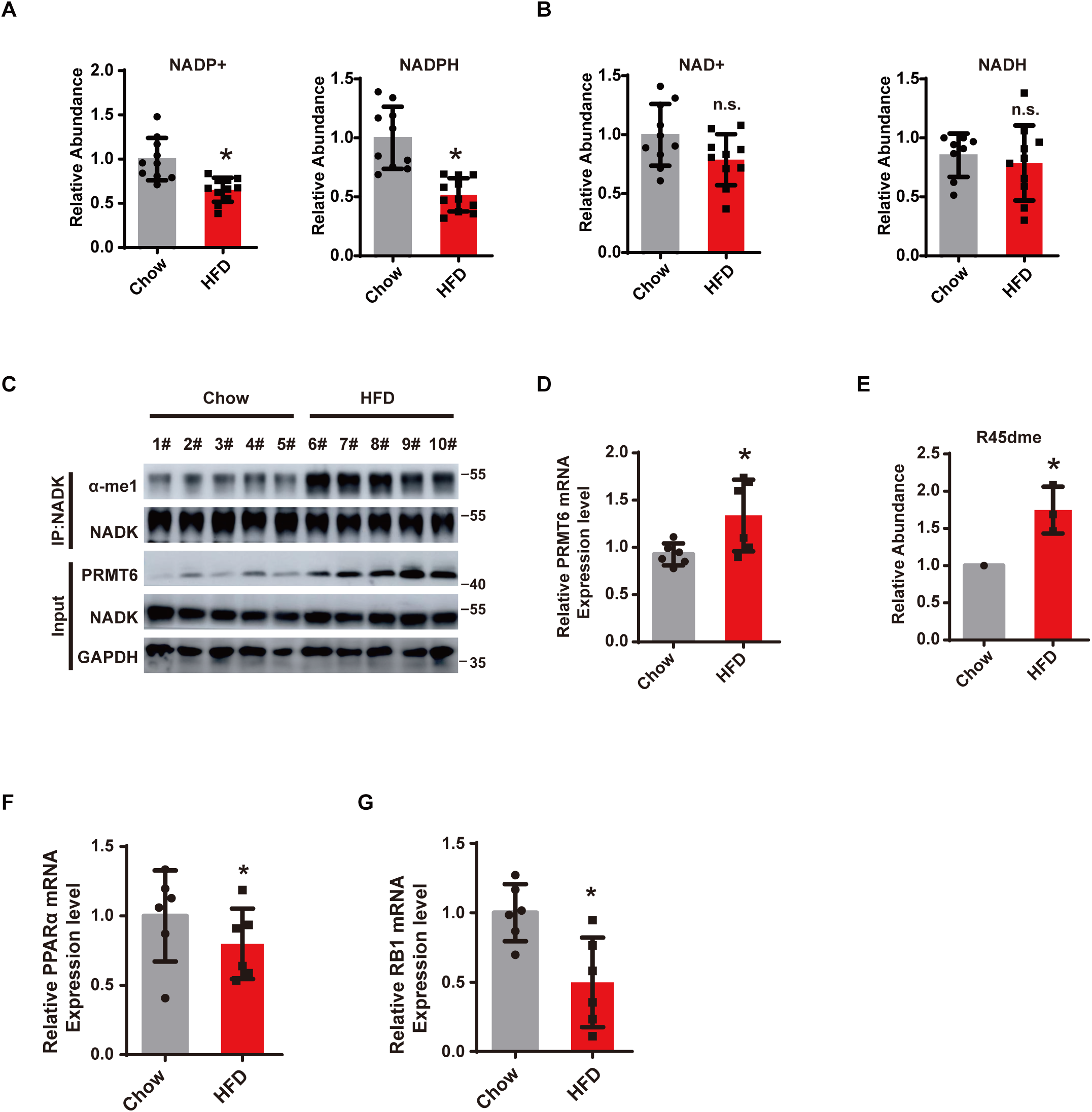
Upregulation of the PRMT6-NADK axis inhibits NADP^+^ synthesis in high-fat diet mice. (A-B) LC-MS quantification of NADP (H) (A) and NAD (H) (B) in the liver of C57BL-6 mice fed with chow or 60% HFD for 18 weeks. (C-E) Western blot analysis for PRMT6 protein levels and NADK methylation levels in the liver of chow or 60% HFD fed mice (C). mRNA levels of PRMT6 (D) and LC-MS/MS quantification of R45 di-methylation on NADK from the liver of chow or 60% HFD fed mice (E). (F-G) mRNA levels of PPARα (e) and RB1 (F) in the liver of chow or HFD fed mice. [(A), (B)] Data are presented as the mean ± SD (n=9). [(D), (F) and (G)] Data are presented as the mean ± SD (n=6). *P < 0.05 for pairwise comparisons calculated using a two-tailed Student’s t test.

Furthermore, we intend to explore how PRMT6 is upregulated by high-fat diet. PRMT6 levels can be regulated by RB1/E2F pathway, and RB1 is positively regulated by the lipid sensor PPARα36. In our HFD mice model, PPARα mRNA level was lower than that in the chow-fed mice (Fig. 6E), and it led to a lower RB1 mRNA level (Fig. 6F). Rb1 protein binding inhibits the transcriptional activation ability of E2F factors^37, 38^. So more E2F bound to PRMT6 promoter to increase PRMT6 mRNA level (Fig. 6C). Taken together, our findings demonstrate that PRMT6 is upregulated by RB1/E2F pathway to suppress NADP^+^ production in high-fat diet mice (Fig. 7).

**Figure 7.**
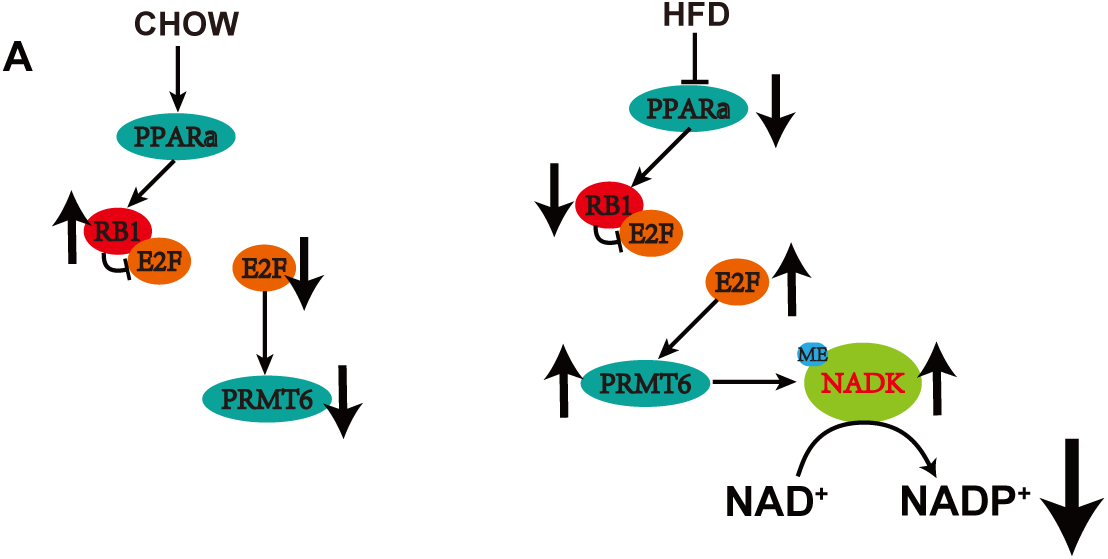
A working model for NADK regulation by the PRMT6-mediated methylation pathway. (A) In the chow-fed condition, RB1 is not inhibited, resulting in suppression of E2F, which leads to a decrease in PRMT6 level. This leads to a low NADK methylation level and high activity, promoting NADP^+^ synthesis. (B) In HFD-fed mice, the lipid sensor PPARα is inhibited, leading to a reduction in RB1 level. This causes an increase in E2F, promoting PRMT6 level, and subsequently attenuating NADP^+^ synthesis through NADK methylation.

## Discussion

Given the dynamic flexibility and multiple roles of NAD^+^/NADP^+^ in cells ^1,3^, it is important to understand the molecular basis for how NAD^+^/NADP^+^ homeostasis contributes to cellular function and how they are regulated by various physiological or pathological conditions. Here we reveal that NADK, the key enzyme of NADP^+^ synthesis from NAD^+^, is inhibited by PRMT6-mediated arginine methylation. PRMT6-mediated methylation of NADK attenuates NADP^+^ synthesis in an Akt-dependent and -independent manner. Furthermore, PRMT6-NADK axis is associated with obesity using a high-fat diet mouse model. Downregulation of NADK methylation in HCC enhances NADP^+^ production to promote cancer development. Our data highlight the highly complex nature of the biological effects of obesity and energy status, which involves reprogramming cellular metabolism, such as NADP^+^ synthesis and homeostasis.

Both our data and previous studies suggest that the N-terminal of NADK is the major regulatory region for NADK activity^14,19,20,36^. Structural studies on NADK (PDB code: 3PFN) show that the N-terminal is mainly flexible, and natural disordered region prediction using several algorithms consistently indicates that the N-terminal is disordered. The N-terminal is outside the NADK enzymatic domain, but it could sense different environmental cues directly or indirectly, such as insulin signaling and high fat. Insulin-induced Akt-mediated phosphorylation of NADK stimulates its activity to increase NADP^+^ production through relief of an autoinhibitory conformation inherent to its amino terminus. Lipid storage inhibits NADP^+^ synthesis through inducing PRMT6-mediated methylation of NADK. NADK methylation enhances this inherent auto-inhibitory conformation. As two different post-translational modifications, phosphorylation and methylation happen in the same regulatory N-terminal region of NADK and antagonize each other (Fig. 5A, B). However, methylation-mediated NADK inhibition is in a phosphorylation-independent manner. In obesity, NADK phosphorylation is inhibited and methylation is elevated, both down-regulating NADK activity. More structural or biophysical studies are required to understand how the N-terminal affects NADK activity.

Interestingly, a previous study showed that high-fat diet inhibited the PI3K/Akt signaling pathway in the fatty liver^37^. A more recent study discovered that PRMT6 methylates PTEN at R159 to suppress PI3K-Akt Signaling^38^. Our data, along with the above results, imply that PRMT6 not only promotes NADK methylation to attenuate NADK activity, but also inhibits NADK activity by reducing Akt-mediated NADK phosphorylation. Previous studies have also uncovered the molecular mechanisms by which PRMT regulates cellular transformation^17^. Thus, our results illustrate a novel mode of PRMT6-mediated NADK methylation influencing cell redox and growth.

In obesity or the elderly population, lower lipid sensor PPARα leads to a lower RB1 level, which in turn causes a higher level of E2F to increase PRMT6 level, thus inhibiting NADK activity via N-terminal methylation (Fig. 5C-F). It is not surprising that the PRMT6-NADK axis is linked to obesity, type 2 diabetes, and cancers, because NADP^+^ is fundamental for anabolic reactions and redox balance. Our studies suggest that controlling the activity of NADK could be a new strategy for NADPH-associated diseases, such as cancer and obesity.

## Materials and Methods

### Antibodies and other reagents

Antibodies to Akt (Cell Signaling Technology (CST), 4691), NADK (CST, 55948), p-RRXpS/pT (CST, 9624), PRMT6 (CST, 14641), Mono-Methyl Arginine [α-mme-R] (CST, 8015), Asymmetric Di-Methyl Arginine Motif [α-me2a-R] (CST, 13522), horse radish peroxidase (HRP)-conjugated secondary antibodies anti-mouse (CST, 7076) and anti-rabbit (CST, 7074) were purchased from CST, anti-FLAG-M2 (Sigma, F1804), anti-HA (Sigma, H3663), and anti-β-actin (Sigma, A5316) were purchased from Sigma. Another NADK antibody was from Santa Cruz Biotechnology, sc-100347. anti-FLAG M2 Affinity Gel (A2220) was from Sigma.

#### Other reagents including

NAMPT inhibitor FK-866 hydrochloride (Tocris, 480810), β-nicotinamide adenine dinucleotide hydrate (NAD) (Sigma, N7004), D-glucose 6-phosphate disodium salt hydrate (G6P) (Sigma, G7250), glucose-6-phosphate dehydrogenase (G6PD) from baker’s yeast (Sigma, G6378), adenosine triphosphate salt (ATP) (Sigma, A2383), β-nicotinamide adenine dinucleotide phosphate hydrate (NADP) (Sigma, N5755), β-nicotinamide adenine dinucleotide phosphate, reduced tetra (cyclohexylammonium) salt (NADPH) (Sigma, N5130), folic acid (Sigma, F7876), pyridoxal hydrochloride (Sigma, P6155), riboflavin (Sigma, R9504), protease inhibitor cocktail (Sigma, P8340), IPTG (Sigma, I6758), ^13^C3-^15^N-nicotinamide (Cambridge Isotope Labs, CNLM-9757-0.001), Lipofectamine RNAiMAX (ThermoFisher Scientific, 13778), Lipofectamine 2000 Transfection Reagent (Thermo Fischer, 11668), HEPES (Thermo, 15630-080), Glutathione (Thermo, 78259), sodium pyruvate (Thermo, 11360-070), Corning® glutagro™ (Corning, 25-015-CI), Gibco™ 2-mercaptoethanol (Gibco, 21985023), BCA kit (Thermo, 23225), hygromycin B (Life Technologies/Invitrogen, 10687-010), polyethylenimine (PEI) (Polysciences, 23966-2), nicotinamide-free DMEM (US biologicals, D9800-17), thiamine hydrochloride (Santa Cruz, sc-205859), polybrene (Santa Cruz, sc-134220), KOD-Plus-Neo (Toyobo, KOD-401), Lonza SeaPlaque® Agarose (VWR, 50101), and dialyzed serum (dFBS; Life Technologies/Gibco, 26400-036) were used as indicated.

#### Cell culture

HEK293T (ATCC), and HepG2 (ATCC) cell lines were cultured in Dulbecco’s Modified Eagle Media (DMEM) supplemented with 10% FBS and 1% Penicillin-Streptomycin (P/S) in a 37°C incubator with a humidified, 5% CO2 atmosphere.

#### Animal Studies

Four-week-old male C57BL/6 mice (n=40) were purchased from SiBeiFu (Beijing) and housed in the Chinese National Center for Protein Science animal breeding facility. The C57BL/6 mice were housed in a temperature-controlled environment under a standard light/dark cycle (12 h/12 h) and were given ad libitum feeding. The Animal Care and Use Committee of the National Center for Protein Science approved all the mice experiments. Half of the group were fed with a standard “Chow” diet containing 13.7 kJ g^-1^, and the other half were fed with a 60% HFD containing 21.8 kJ g^-1^. Body weight were measured weekly, and both groups were sacrificed at 11 weeks of treatment. Mice were sedated using isoflurane and sacrificed by cardiac puncture. The liver was separated, and one part of the liver was fixed in 4% PFA solution. Another part was snap-frozen in liquid nitrogen for LC-MS based metabolite profiling, H&E staining, or Western blot analysis, and the remaining part of the liver was quickly cut into smaller pieces and snap-frozen in liquid nitrogen. Samples were stored at -80°C and pulverized in liquid nitrogen prior to analysis.

#### cDNA constructs, RNAi, and CRISPR/Cas9

Full-length human NADK isoform 1 was cloned into the pIRES2-S-SBP-FLAG, pGEX-4T-2, FUGW, pHAGE and pET-28a vectors. NADK variants R39K, R41K, R45K, R39/R45K, R39/R41/R45K, S44/46A were generated by PCR-based site-directed mutagenesis using KOD-Plus-Neo DNA Polymerase, with mutations verified by sequencing. HA-Akt-CA (myristoylated-Akt) and HA-PRMT6 were constructed into the pCMV vector.

HEK293T cells expressing non-targeting control shRNA (pLKO.1-puro) or shRNA targeting NADK or PRMT6 were generated via lentivirus delivery and puromycin (2 μg ml^−1^) selection. The resulting stable cell lines were used to generate stable NADK-reconstituted cell lines using lentiviral NADK expression vectors (pHAGE-Hygro) following hygromycin (300 μg ml^−1^) selection.

All siRNA oligonucleotides were purchased from Sigma-Aldrich. Cells were transfected twice at a 24-hour interval with the indicated siRNA using Lipofectamine® RNAiMAX (Invitrogen) according to the manufacturer’s instructions. The siRNA target sequences are in the resource table.

PRMT 6-1: CACCGGCATTCTGAGCATCTT

PRMT 6-2: ACCAGTGGAGACTGTAGAGTT

NADK KO HEK293T and HepG2 cells were generated via CRISPR/CAS9 method using LentiCRISPR v2 system. The guide RNA used to generate NADK KO cells was: 5’-AACTCCAGGTCTCATCGCCG-3’. Cells integrated with guide RNA and Cas9 were selected with 2 μg ml^−1^ puromycin, and then single clones were picked.

#### Immunoprecipitation and Immunoblotting

Cells transfected with indicated constructs were collected and lysed in NETN buffer (10 mM Tris-HCl pH 8.0, 100 mM NaCl, 1 mM EDTA, and 0.5% NP-40) supplied with protease inhibitors (Roche) on ice for 30 min. Then cell lysates were subjected to indicated beads (For flag-tagged protein immunoprecipitation, FLAG M2 (Sigma) were used. For HA-tagged protein immunoprecipitation, HA beads (Sigma) were used). After 8 h rotation at 4°C, beads were washed with NETN buffer three times, and samples were eluted with 2×SDS loading buffer for immunoblotting with indicated antibodies.

For endogenous IP, cell lysates were incubated with indicated antibodies at 4°C for 6 h, and then were subjected to Protein A/G beads (Thermo fisher) for 4 h at 4°C. Beads were then washed with NETN buffer three times, and samples were eluted with 2×SDS loading buffer for immunoblotting with indicated antibodies.

For Western blotting, proteins were transferred to nitrocellulose membrane after SDS-PAGE using the standard procedures. The nitrocellulose membrane was blocked with 5% BSA in TBST at room temperature for 1 h, and the excess reagents were removed by repeated washing using TBST. The nitrocellulose membrane was incubated with antibodies (1:1000 dilution) in 0.5% BSA in TBST overnight at 4°C. Non-specifically adsorbed antibodies were removed by washing the nitrocellulose membrane three times with TBST. The nitrocellulose membrane was incubated with goat anti-rabbit or anti-mouse IgG-HRP conjugate (1:2000 dilution) in 0.5% BSA in TBST at room temperature for 1 h. Finally, after washing three times with TBST, the nitrocellulose membrane was stained with luminol and peroxide detection solution (Thermo Fisher Scientific, USA) at 1:1 ratio in the dark for 1 min and imaged using a GE ImageQuant LAS 500 system (GE Healthcare, Chicago, IL, USA).

Recombinant protein purification: GST fusion proteins were expressed and purified from *E.coli* BL21 (DE3) strain, and were immobilized on Glutathione Sepharose 4B (GE healthcare) at 4°C overnight. Proteins were then eluted with 5-fold volume of 10 mM Glutathione solution, and were concentrated using 10 kD cutoff protein concentrators (Thermo fisher). Proteins were aliquoted and stocked in PBS containing 5% Glycerol at -80°C.

#### GST pull-down Assay

GST fusion proteins were immobilized on Glutathione Sepharose 4B at 4°C overnight. HEK293T cells transfected with indicated constructs were lysed in NETN buffer supplied with protease inhibitors, and were incubated with the Sepharose immobilized with indicated proteins at 4°C for 8 h. Sepharose was then washed with NETN buffer three times and boiled in 60 μl of 2×SDS loading buffer for subsequent immunoblotting with indicated antibodies.

#### *In vitro* methylation assay

For *in vitro* methylation assay, GST-tagged PRMT1 or PRMT6 was mixed with recombinant substrate GST-NADK in methylation reaction buffer (50 mM Tris–HCl, pH 8.0, 20 mM KCl, 5 mM DTT, 4 mM EDTA) in the presence of 200 μM S-adenosyl-L-methionine (SAM) (Sigma) and incubated at 37°C for 1 h in a final volume of 30 μl. Reactions were stopped by adding SDS-PAGE loading buffer, followed by boiling for 10 min, and subjected to western blotting analysis.

Real-time PCR analysis: Total RNA was extracted using an RNA extraction kit from Beijing BLKW Biotechnology following the manufacturer’s instructions. 1 μg RNA was used for reverse transcription by FastQuant room temperature Kit (TIANGEN) according to the manufacturer’s protocol. Real-time PCR was performed using SYBR green (Bio-Rad), and the results were analyzed by a Bio-Rad CFX96 machine. All qPCR assays were repeated three times, and the primers were included in the resource table.

PRMT6 Human Forward: TACCGCCTGGGTATCCTTCG

PRMT6 Human Reverse: CCTGTTCCGGCAACTCTACA

PRMT6 Mouse Forward: TACTACGAGTGCTACTCCGAC

PRMT6 Mouse Reverse: CTAAGCGGTAGGCTTCGGT

PPARα Mouse Forward: TACTGCCGTTTTCACAAGTGC

PPARα Mouse Reverse: AGGTCGTGTTCACAGGTAAGA

RB1 Mouse Forward: TCGATACCAGTACCAAGGTTGA

RB1 Mouse Reverse: ACACGTCCGTTCTAATTTGCTG

NADK Human Forward: CGACGGAGTGATCGTGTCC

NADK Human Reverse: GTGTTCCTTGCTTCAGGTGAC

RB1 Mouse Forward: TGAGACCTGGAGCTACAATCA

RB1 Mouse Reverse: ATGACAAGCACACTCTTGGGA

#### Sequence Alignments

NADK amino acid sequences from different species were obtained from Uniprot (http://www.uniprot.org/). Multiple sequence alignments were done using Clustal Omega. The alignments were visualized in Jalview (http://www.jalview.org/).

#### NADK enzymatic assay

Approximately 0.5 μg of purified NADK or NADK variants were subjected to an NADK enzymatic assay that couples its generation of NADP^+^ to G6PD-mediated production of NADPH, which was then measured as a change in absorbance at 340 nm (A340) over time. The assay was performed in a 96-well plate containing 100 μl of reaction buffer: 10 mM ATP, 10 mM glucose-6-phosphate, 0.5 U G6PD, 10 mM MgCl2, 100 mM Tris-HCl (pH 8.0), and varying concentrations of the substrate NAD^+^ (1 mM, 2 mM, 5 mM, 10 mM). Measurements of A340 were acquired every 2 min for 20 min at 37°C. Standard curves were generated from pure NADPH standards, experimental data were imported into GraphPad Prism 8 to determine the Vmax and KM values. Kcat=Vmax/[E].

LC-MS based metabolite profiling and ^13^C3-^15^N-nicotinamide tracing: For ^13^C3-^15^N-nicotinamide tracing studies, cells were washed once with serum- and nicotinamide-free DMEM. Subsequently, they were incubated in the same medium containing 4 mg L^-1^ of ^13^C3-^15^N-nicotinamide for 1 h, as indicated. In experiments involving inhibitors and growth factors, cells were pretreated with inhibitors for 30 min prior to the co-addition of growth factors in nicotinamide-free DMEM containing ^13^C3-^15^N-nicotinamide. The nicotinamide-free media containing labeled nicotinamide was prepared using powdered DMEM (USBiological Life Sciences, D9815) (Hoxhaj et al., 2019), supplemented with standard DMEM concentrations of Folic Acid, Nicotinamide, Pyridoxal, Riboflavin, Thiamine, Glucose, and Sodium Pyruvate. Nicotinamide was replaced with ^13^C3-^15^N-nicotinamide, and the pH was adjusted to 7.2.

The DMEM medium was discarded and the cells were rapidly washed three times with chilled PBS. The cells were scraped in PBS and then were transferred to 1.5 mL Eppendorf tubes. The cell suspensions were centrifuged at 1000×g for 3 min at 4°C to remove PBS and the cell pellets were shock-frozen in liquid nitrogen, and then 500 µl of MeOH/H2O (80:20) was added for cell disruption and protein precipitation. The resulting solution was then incubated in a rotary shaker at 1000 rpm for 3 min at 80°C. The cell lysate/methanol mixture were centrifuged at 4°C for 10 min at 13,200×g to remove the protein pellet. The supernatants were freeze-dried for LC-MS analysis. For all extractions, the remaining pellets were resuspended in 8 M Urea, 10 mM Tris, pH 8, heated to 60°C for 1 h with shaking, then after centrifugation, the protein concentration in the supernatant was quantified using BCA kit.

The ^13^C3-^15^N-nicotinamide tracing studies were performed using a ACQUITY UPLC H-Class system coupled with a 6500 plus QTrap mass spectrometer (AB SCIEX, USA) equipped with a heated electrospray ionization (HESI) probe. Extracts were separated by a Synergi Hydro-RP column (2.0×100 mm, 2.5 μm, Phenomenex) with a flow rate of 0.5 mL min^-1^. Eluent A was 2 mM triisobutylamine adjusted with 5 mM acetic acid in water, and eluent B was methanol. Separation was operated under the following linear or gradient conditions: 0∼1 min at 5% B; 1∼6 min, 5∼50% B; 6∼7 min, 98% B; 7∼8 min, 5% B. Column chamber and sample tray were held at 35°C and 10°C.

Metabolites were detected in multiple reaction monitor (MRM) mode. MS conditions were: nebulizer gas (Gas1), heater gas (Gas2), and curtain gas were set at 55, 55, and 35 psi, respectively. The ion spray voltage was - 4500 V in negative ion mode. The optimal probe temperature was determined to be 500°C. The SCIEX OS 1.6 software was applied for metabolite identification and peak integration.

LC-MS/MS-based mono-methylated/dimethylated-peptide analyses: NADK purified from HEK293T cells was digested with Glc-C and Lys-C mixture (sequencing grade, Promega, protein: Glc-C: Lys-C = 20:1:1) at 37°C for 16 h and desalted using C18 Zip-Tips (Millipore).

LC-MS/MS analysis was carried out using an Easy nLC-1200 system coupled with an Orbitrap Fusion Lumos Tribrid mass spectrometer (Thermo Fisher Scientific). The peptides were resuspended using 10 µl of 0.1% FA aqueous solution and were separated using in-house-made 15 cm length columns (150 mm i.d,) packed with Ultimate XB-C18 1.9 µm reverse phase resin (Welch materials). A binary solvent system consisting of buffer A (0.1% formic acid in water) and buffer B (0.1% formic acid in acetonitrile) was employed for chromatographic separation with a constant flow rate of 600 nL min^-1^. The gradient was set as follows: 7%∼15% B for 11 min, 15%∼25% B for 37 min, 25%∼40% B for 20 min, 40%∼100% B for 1 min, and 100% B for 6 min. The mass spectrometry parameters were set as: spray voltage of 2 kV and ion transfer tube temperature of 300°C. The eluted peptides were analyzed by data-dependent acquisition (DDA) with a dynamic exclusion duration of 18 s. The m/z scan range was 350 to 1550 for full scan, and the intact peptides were detected in the Orbitrap at a resolution of 60,000. Parameters for peptide fragmentation were set as: type 1 was EThcD mode with a supplemental activation (SA) collision energy of 25%, type 2 was HCD mode with collision energy of 30%. The peptide fragments were detected in the Orbitrap at a resolution of 30,000 with AGC value of 4e5 and maximum injection time of 50 ms.

The mass spectrometric raw data were searched using MaxQuant software (version 1.6.1.0) against the UniProt NADK protein sequence (O95544). The mass tolerances were 20 ppm for precursor ions and 10 ppm for fragment ions. Two missed cleavages were allowed and Carbamidomethylation **c** was set as a fixed modification, while oxidation (M), acetylation (protein N-term), and Methylation/Dimethylation (R) were set as variable modifications. The false discovery rate (FDR) cut-off for peptide-spectrum matches (PSMs) and proteins identification was set at < 1%.

Absolute quantification of peptide NADK arginine methylation was performed using a Q Exactive™ mass spectrometer (MS, ThermoFisher Scientific, USA) connected to a Dionex UltiMate 3000 RS UPLC system. MS analysis was operated in parallel reaction monitoring mode for peptide and NADK digestion quantification. Targeted proteomic data was processed by Skyline (version 23.1).

### NADK peptides

TWSYNHPIRGRAK; SRSLSASPALGSTK; TWSYNHPIR(me1)GRAK; TWSYNHPIR(me2)GRAK; TWSYNHPIRGR(me1)AK; TWSYNHPIRGR(me2)AK; SR(me1)SLSASPALGSTK; SR(me2)SLSASPALGSTK; Colony formation assays: Colony formation assays were performed as described previously (Pei et al., 2011).

Xenograft tumor growth assays: Three-week-old male nude mice (n = 40) were purchased from Vital River (Beijing) and housed in Chinese National Center for Protein Science animal breeding facility. A total of 1x10^6^ cells were inoculated into 8-week-old male nude mice. After the tumor size reached 2.5 mm in diameter, the tumor size and body weight of each group were measured every week. At the indicated times, the mice were euthanized, and the tumors were weighed.

Isothermal Titration Calorimetry (ITC) assay: The ITC experiment was conducted on the MicroCal PEAQ-ITC. After cleaning the sample cell and syringe, the unmethylated and methylated NADK proteins was carefully injected into the sample cell using a bubble-free micro-injector. NAD^+^ or ATP was filled into a 40 μL titration syringe. All proteins and NAD^+^ or ATP were prepared in Hepes buffer (20 mM Hepes-NaOH, 150 mM NaCl, pH 7.0), and the reference cell were injected with deionized water as a heat-balancing control. After titrating the sample cell 19 times with NAD^+^ or ATP at a constant rate of 150 s, the titration curve was fitted using the nonlinear least squares estimation model in MicroCal PEAQ-ITC Analysis Software (Version 1.41).

## Data Availability

The authors declare that all data supporting the findings of this study are available within the paper and its Supplementary Information files. Further information and requests for resources and reagents should be directed to W.Q., S.H., Y.C. and P.Z.

## Supplementary Materials

**Figure S1.**
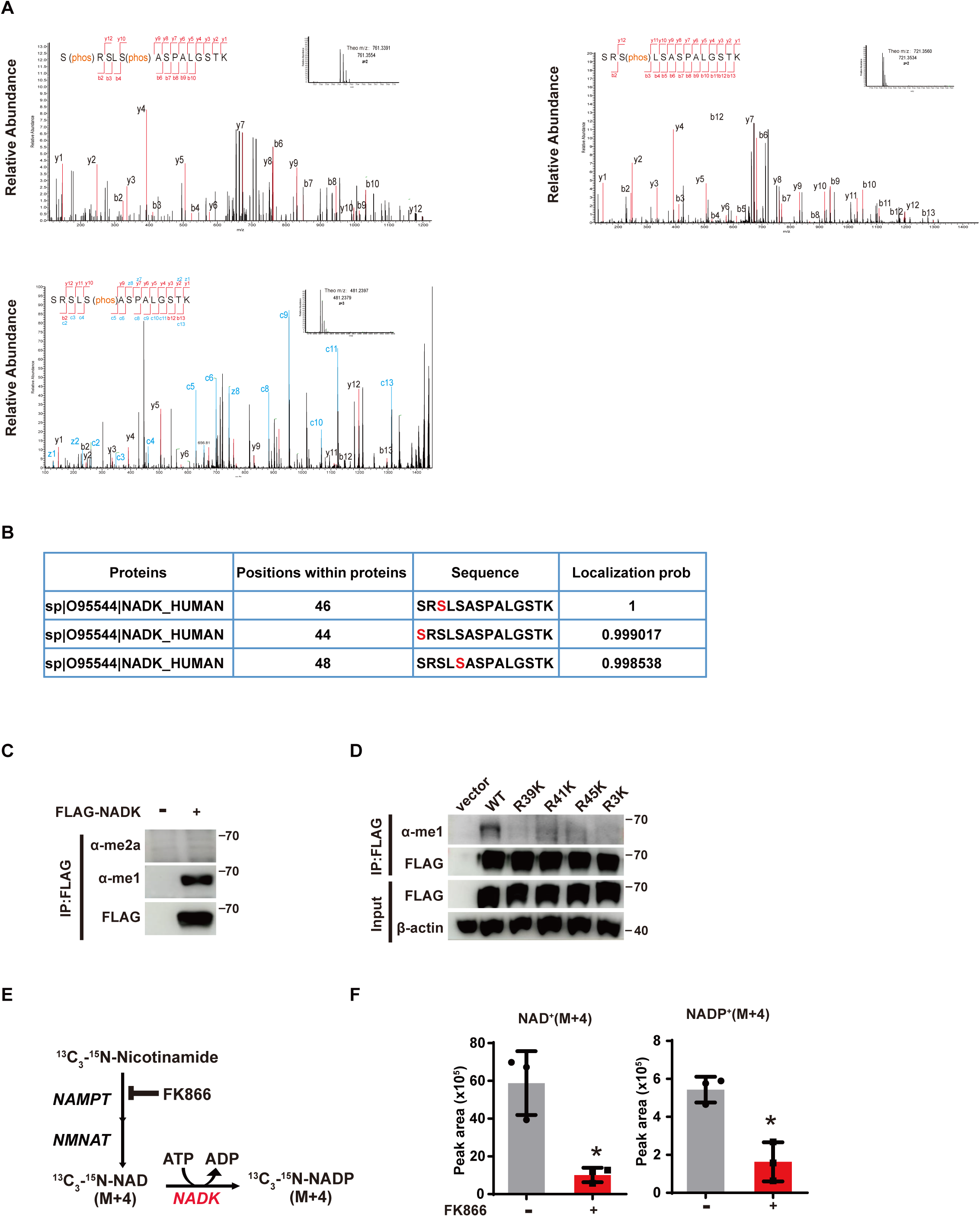
NADK N-terminal is methylated. (A) Representative MS/MS spectrum for peptides containing phosphorylation on residues Ser44, Ser46, and Ser48. Endogenous NADK was immunopurified from HEK293T cells and analyzed by LC-MS/MS as described in the methods. (B) As in (A), the phosphorylation sites are characterized and evaluated. (C) FLAG-NADK expressed in HEK293T cells were immunoprecipitated and immunoblotted with anti-Mono-Methyl Arginine or anti-Di-Methyl Arginine antibody. (D) FLAG-NADK WT and variants were transfected into HEK293T cells, and NADK methylation was detected with anti-Mono-Methyl Arginine antibody. (E) Schematic of labeling. Salvage enzyme NAMPT (nicotinamide phosphoribosyl-transferase) is selectively inhibited by FK866 to block M+4 derivatives of NAD^+^ and NADP^+^ synthesized from the salvage pathway. (F) *De novo* synthesized NAD^+^ and NADP^+^ when cells were treated with NAMPT inhibitor FK866. [(F)] Data are presented as the mean ± SD of biological triplicates and are representative of at least three independent experiments. *P < 0.05 for pairwise comparisons calculated using a two-tailed Student’s t test.

**Figure S2.**
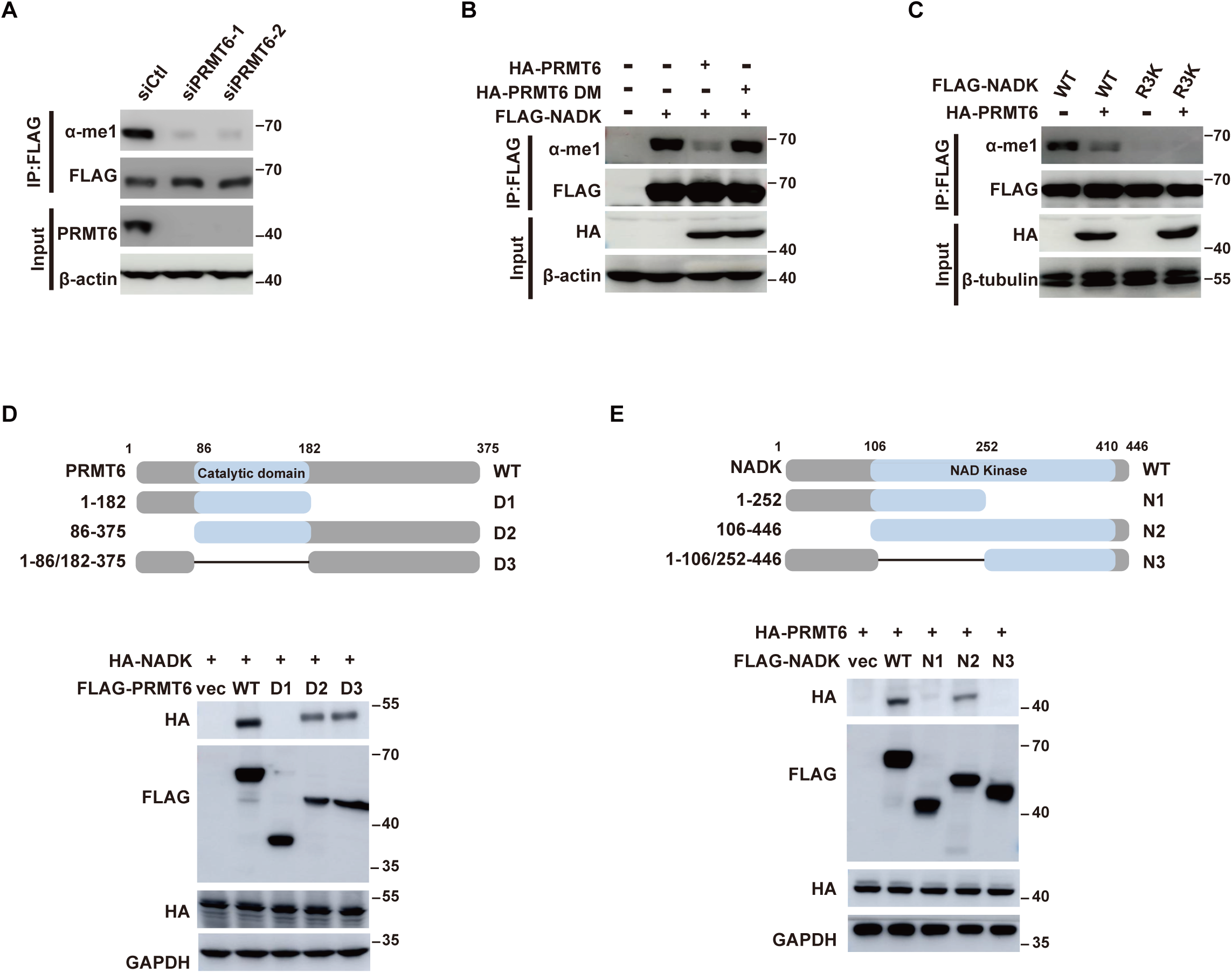
PRMT6 mediates NADK N-terminal methylation. (A) NADK methylation in PRMT6-depleted HEK293T cells was detected with anti-Mono-Methyl Arginine antibody. (B) HEK293T cells were co-transfected with FLAG-NADK and PRMT6 or PRMT6-DM (V86A/L87A/D88A mutant). Methylation on FLAG immunoprecipitated NADK was detected as in (A). (C) HA-PRMT6 was co-transfected with FLAG-NADK WT or R3K mutant into HEK293T cells. NADK methylation were detected as in (A). (D) Schematic representation of truncated PRMT6 mutants used in this study. Plasmids encoding FLAG-tagged full-length or truncation/deletion mutants of PRMT6 were co-transfected with plasmids encoding HA-tagged full-length NADK into HEK293T cells. Immunoprecipitation and immunoblotting were performed 48 h post transfection. (E) Schematic representation of NADK truncation mutants used in this study. Plasmids encoding FLAG-tagged full-length or truncation mutants of NADK were co-transfected with plasmids encoding HA-tagged full-length PRMT6 into HEK293T cells. Immunoprecipitation and immunoblotting were performed 48 h post transfection.

**Figure S3.**
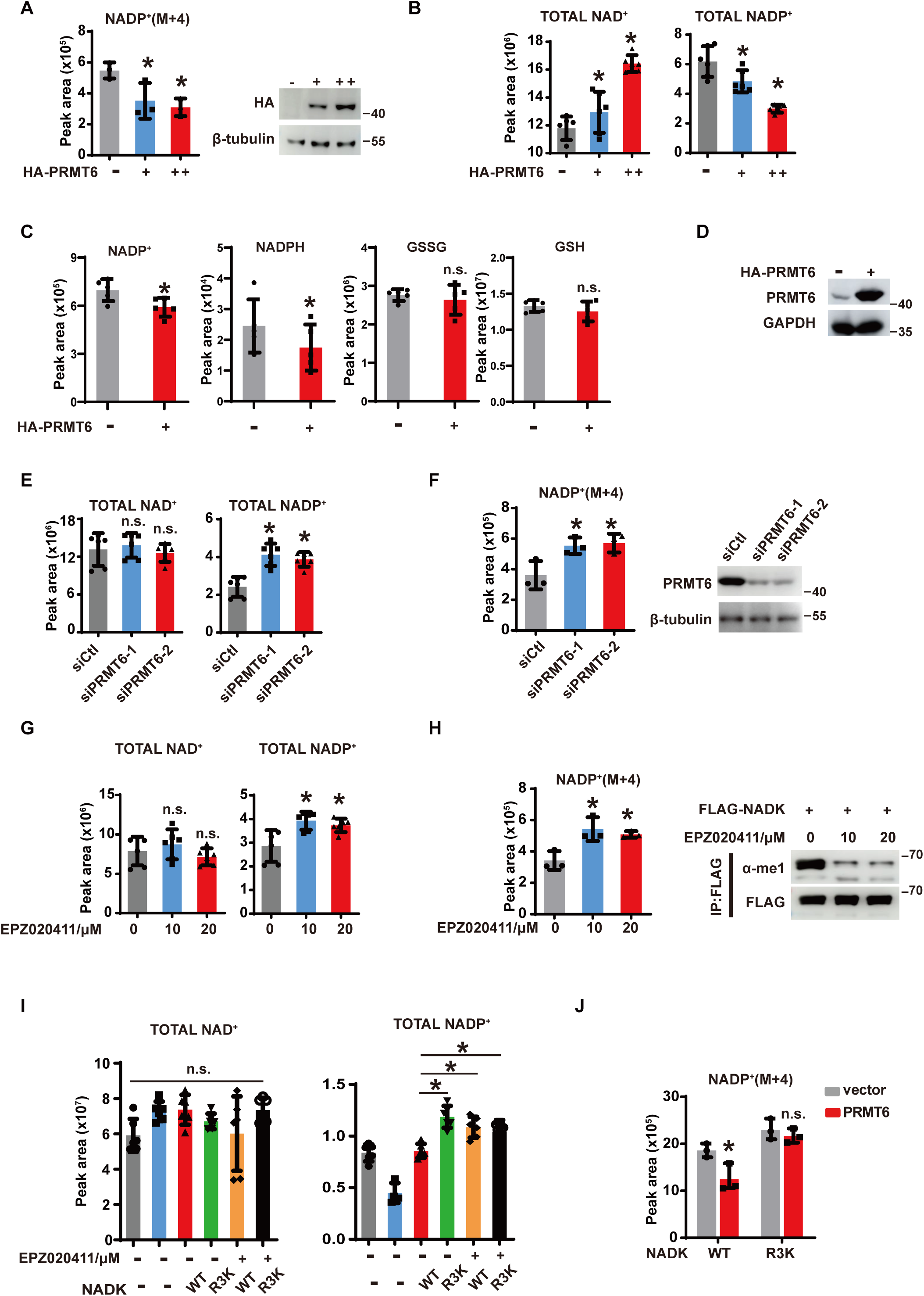
PRMT6-mediated NADK methylation attenuates its kinase activity. (A) Quantification of *de novo* NADP^+^ synthesis by LC-MS. HEK293T cells were transfected with HA-PRMT6 plasmids (dose dependent). PRMT6 protein levels were detected by western blot with indicated antibodies. (B) Quantification of total NAD^+^ and NADP^+^ by LC-MS. HepG2 cells were transfected with HA-PRMT6 plasmids (dose dependent). (C-D) Quantification of NADP^+^, NADPH, GSSG, and GSH by LC-MS. HepG2 cells were transfected with HA-PRMT6 plasmids (C). PRMT6 protein levels were detected by western blot with indicated antibodies (D). (E) As in (A), but HepG2 cells were transfected with PRMT6 siRNA, and the total NAD^+^ and NADP^+^ was quantified by LC-MS. (F) As in (B), but HEK293T cells were transfected with PRMT6 siRNA, and *de novo* NADP^+^ synthesis was quantified by LC-MS. PRMT6 protein levels were detected by western blot with indicated antibodies. (G) As in (A), but HepG2 cells were treated with the PRMT6-specific inhibitor EPZ020411 in a dose gradient to inhibit NADK methylation, and the total NAD^+^ and NADP^+^ was quantified by LC-MS. (H) As in (B), but HEK293T cells were treated with the PRMT6-specific inhibitor EPZ020411 in a dose gradient to inhibit NADK methylation, and *de novo* NADP^+^ synthesis was quantified by LC-MS. NADK methylation was detected with anti-Mono-Methyl Arginine antibody. (I) Quantification of total NAD^+^ and NADP^+^ by LC-MS. NADK-depleted HepG2 cells were reconstituted with NADK WT or R3K mutant and treated with the PRMT6-specific inhibitor EPZ020411. (J) Quantification of *de novo* NADP+ synthesis by LC-MS when PRMT6 and NADK WT or R3K mutant were overexpressed in HEK293T cells. [(A), (F), (H) and (J)] Data are presented as the mean ± SD of biological triplicates and are representative of at least three independent experiments. [(B), (C), (E), (G) and (I)] Data are presented as the mean ± SD (n=6). *P < 0.05 for pairwise comparisons calculated using a two-tailed Student’s t test.

**Figure S4.**
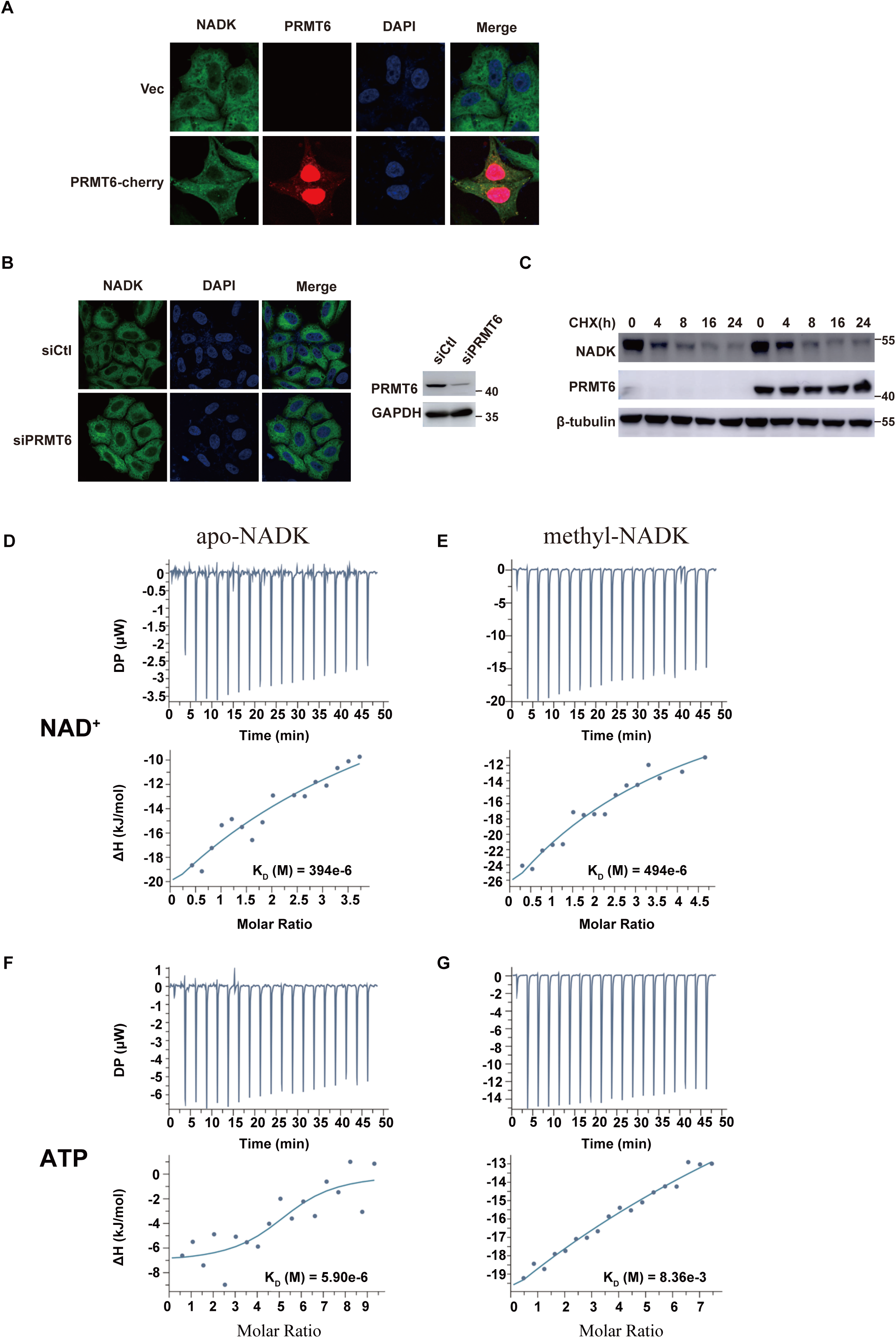
NADK methylation affects ATP binding, but not subcellular localization and stability. (A-B) HepG2 cells were co-transfected with PRMT6-cherry plasmids (A) or siRNAs (B) with NADK-GFP, and NADK cellular localization was detected by GFP fluorescence. PRMT6 protein levels were detected by western blot with indicated antibodies. (C) HEK293T cells infected with PRMT6-HA plasmids were treated with 100 mg ml^-1^ CHX for different time courses. NADK and PRMT6 protein levels were detected by western blot with indicated antibodies. (E-G) The binding affinity (Kd) of NAD^+^ and ATP to unmethylated or *in vitro* methylated NADK proteins measured by ITC. Upper: real-time ITC thermograms; Lower: corresponding integrated heat data and fitted curves.

**Figure S5.**
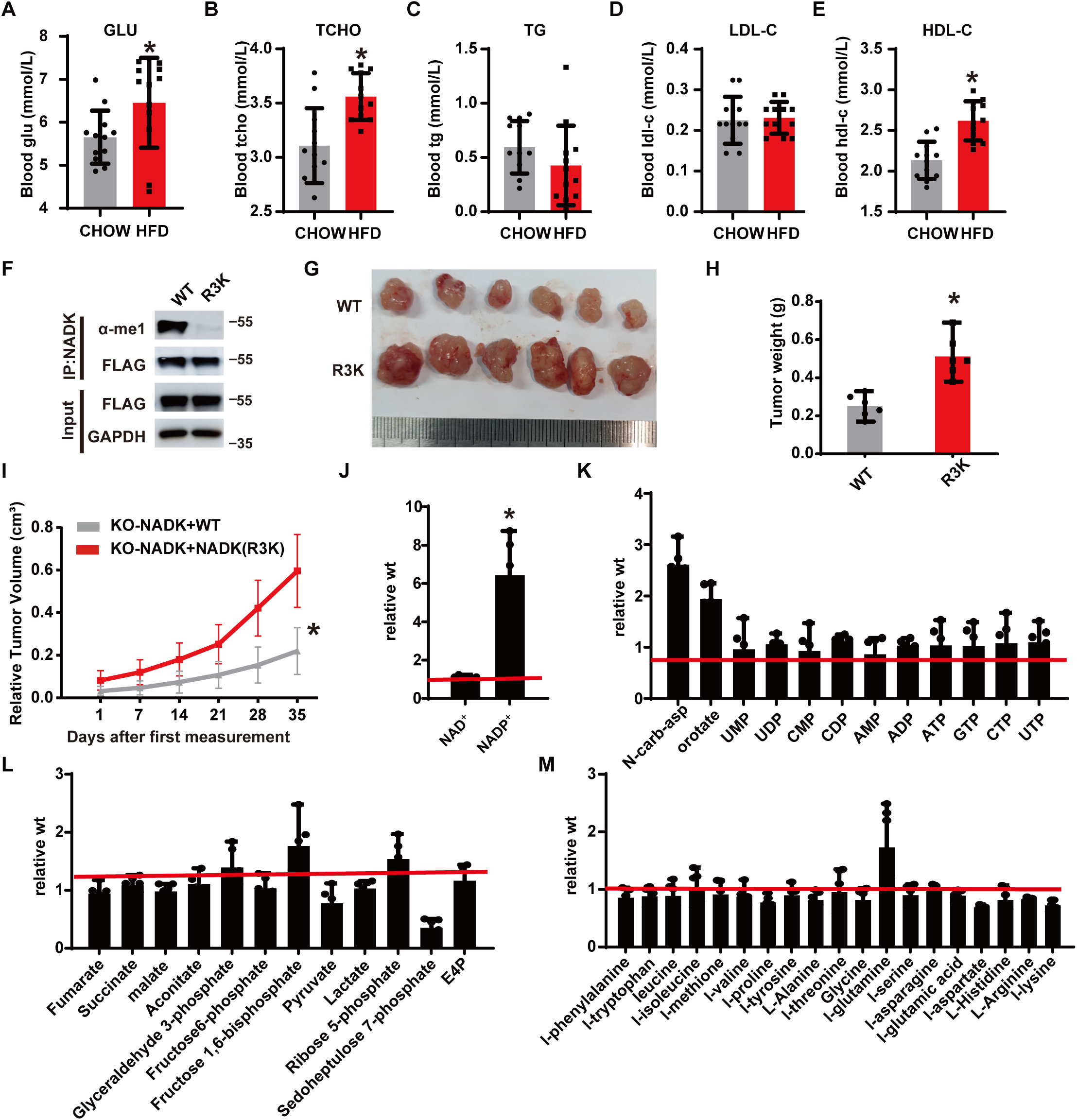
An HFD-induced obese xenografted mice model and down-regulated NADK methylation in liver cancer patients. (A-E) Serum levels of GLU, TCHO, TG, HDL-C, and LDL-C from chow or HFD-fed BALB/c nude mice. (F-I) PRMT6-mediated methylation of NADK suppresses tumor growth in HFD-induced obese xenografted mice. The BALB/c nude mice were fed with 60% HFD for 3 months. Then the NADK WT or R3K reconstituted HepG2 cells were implanted into the obese nude mice by subcutaneous injection. Tumors were harvested by the end of the experiment after mice were sacrificed. NADK methylation was detected with anti-Mono-Methyl Arginine antibody (F). Tumor images (G), tumor weight (H) and tumor volume (I) were presented (n=6). (J-M) NAD^+^ and NADP^+^ levels (J), and nucleotide metabolites (K), glucose metabolites (L), and amino acids metabolites (M) were measured by targeted LC-MS/MS. The indicated metabolites are presented as the NADK R3K tumor abundance in relative to NADK WT tumors, indicated by the dashed red line (n=6). [(A-E)] Data are presented as the mean ± SD (n=10). [(G-M)] Data are presented as the mean ± SD (n=6). *P < 0.05 for pairwise comparisons calculated using a two-tailed Student’s t test.

**Figure S6.**
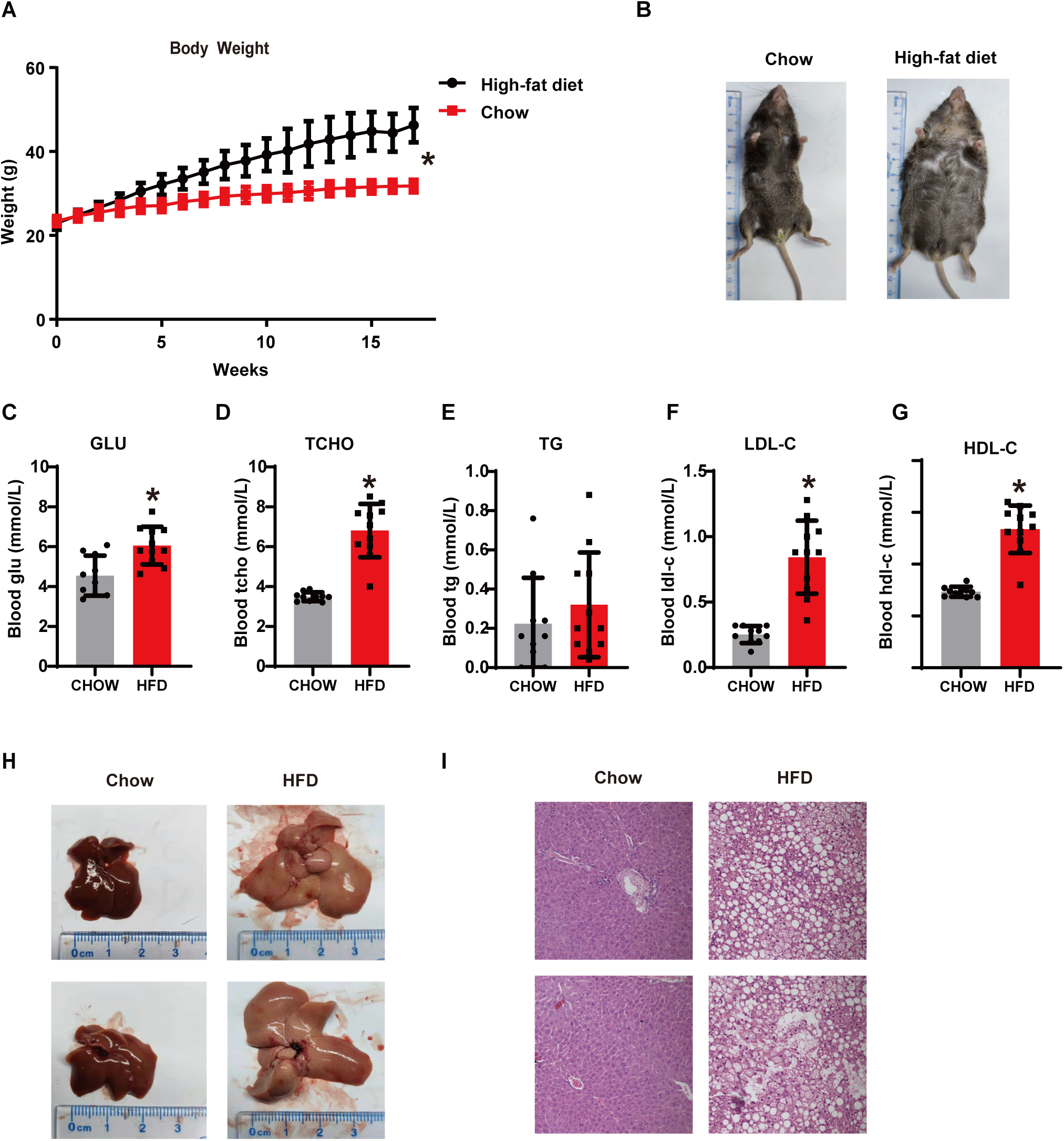
An HFD-induced obese mice model. (A) Total body weight of C57BL-6 mice fed with standard chow or 60% HFD for 4 months (n=20). (B) Representative images from chow or HFD-fed mice. (C-G) Serum levels of GLU, TCHO, TG, HDL-C, and LDL-C from chow or HFD-fed mice. (H) Representative liver images from chow or HFD-fed mice. (I) H&E staining of liver sections from chow or HFD-fed mice. [(A) and (C-G)] Data are presented as the mean ± SD (n=10). *P < 0.05 for pairwise comparisons calculated using a two-tailed Student’s t test.

